# Lipid Composition Modulates Secondary Structure at the Biocondensate-Membrane Interface

**DOI:** 10.64898/2026.05.21.726873

**Authors:** Moonyeon Cho, Carlos R. Baiz

## Abstract

Biomolecular condensates (BMCs) formed through liquid–liquid phase separation (LLPS) play key roles in cellular organization, yet their molecular-scale interactions with membranes remain poorly understood. Here, we use surface-enhanced infrared absorption spectroscopy (SEIRAS) to quantify the secondary structure of poly-GR (glycine–arginine) inside condensates in contact supported lipid bilayers. Amide I SEIRAS provides a surface-sensitive measure of the peptide backbone conformation at the membrane-condensate interface. Results show that neutral POPC bilayers preserve the β-sheet rich organization of the bulk condensate, whereas negatively charged POPC/POPS bilayers reduce β-sheet content, and enhanced β-turn formation, accompanied by perturbations of the lipid headgroups. These results demonstrate that membrane charge modulates condensate secondary structure, hydration, and interfacial behavior, providing molecular-level insight into electrostatic regulation of condensate organization at the membrane-water interface.

**TOC Graphic:** 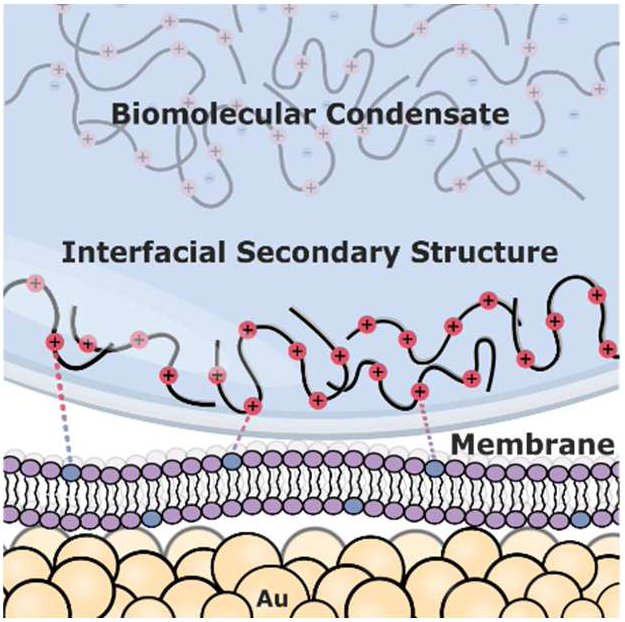

## Introduction

Biomolecular condensates (BMCs) form through liquid–liquid phase separation (LLPS), providing a mechanism for organizing biochemical processes within cells.^1^ Although lacking membranes as physical boundaries, condensates frequently associate with membrane-bound organelles (MBOs) to regulate key processes such as synaptic vesicle clustering,^2^ ribonucleoprotein granule assembly,^3^ and endocytic vesicle trafficking.^4^ The biophysics of condensate-membrane interactions is not well understood. Recent studies have shown that wetting affinity, lipid packing, and interfacial water dynamics influence condensate behavior at membranes,^5–7^ while other work has demonstrated that condensates can remodel membranes. For instance, protein phase separation can induce membrane bending and tubulation,^8–10^ and shift the thermodynamic phase boundaries of lipid bilayers to stabilize membrane domains.^11^ However, while condensate-induced membrane reorganization has been studied,^12^ the reciprocal effect of how specific membrane actively reshape the secondary structure and molecular organization of the proteins within these condensates remains unknown. To understand this, it is important to note that phase separation serves as a critical functional prerequisite; for instance, the linker for activation of T cells (LAT) must assemble into membrane-associated condensates with multivalent partners such as Grb2 and SOS to effectively propagate downstream signaling cascades.^11,13,14^ Within these condensates, proteins exist not as isolated molecules, but as a dense, dynamically cross-linked network. When biomolecular condensates encounter a membrane surface, the resulting structural adaptation is likely governed by the collective physical properties at the condensate-membrane interface. Therefore, to understand the molecular basis of condensate-membrane interactions, it is essential to determine how the condensed phase itself dictates the structural remodeling of proteins at the membrane boundary.

At the molecular level, low-complexity domains (LCDs), which are intrinsically disordered, are prone to condensate formation. Unlike folded proteins with stable secondary structures, IDPs possess high solvent accessibility and fewer stabilizing intramolecular contacts, as such the surrounding environment can directly influence their structural ensembles.^15,16^ These proteins transiently adopt local secondary motifs whose populations shift in response to the environment.^17^ Understanding how these structural ensembles are reshaped by biological interfaces, such as membranes, is a necessary step towards understanding how condensates assemble, maintain their material properties, and interact with surrounding cellular components.

Infrared spectroscopy probes protein secondary structure through amide backbone carbonyl stretching modes, known as the amide I region, where distinct vibrational signatures correspond to different structural motifs.^18–21^ However, characterizing protein structure at membrane-condensate interfaces requires surface sensitivity in addition to spectral specificity. Surface-enhanced infrared absorption spectroscopy (SEIRAS) addresses this challenge by exploiting local electromagnetic field enhancements near nanostructured metal surface, selectively probing the interface with penetration depths below 100 nm (**Figure 1d**). ^22–24^ This probing depth allows SEIRAS to quantify the structure of peptides specifically within the interfacial region of condensate droplets, instead of the bulk droplet. SEIRAS has been widely applied in interfacial studies of electrochemical systems and biological membranes,^22^ yet this work represents its first use in probing interfacial protein structure within biomolecular condensates in contact with lipid membranes. Using SEIRAS, we directly quantify how poly-GR (**Figure 1c**) peptides in condensates are modulated by lipid bilayers with different lipid compositions (**Figure 1a, b**). Poly-GR is derived from the GGGGCC repeat expansion in C9orf72 associated with amyotrophic lateral sclerosis (ALS).^26,27^ In cells, poly-GR localizes to membrane-bound organelles (MBOs), including mitochondria, where it contributes to mitochondrial dysfunction.^28^ Beyond its pathological significance, poly-GR serves as a tractable biophysical model for studying electrostatically driven condensate-membrane systems.^11^ This approach enables molecular-level characterization of protein secondary structure at membrane-condensate interfaces and reveals how membrane composition modulates protein conformation.

**Figure 1.**
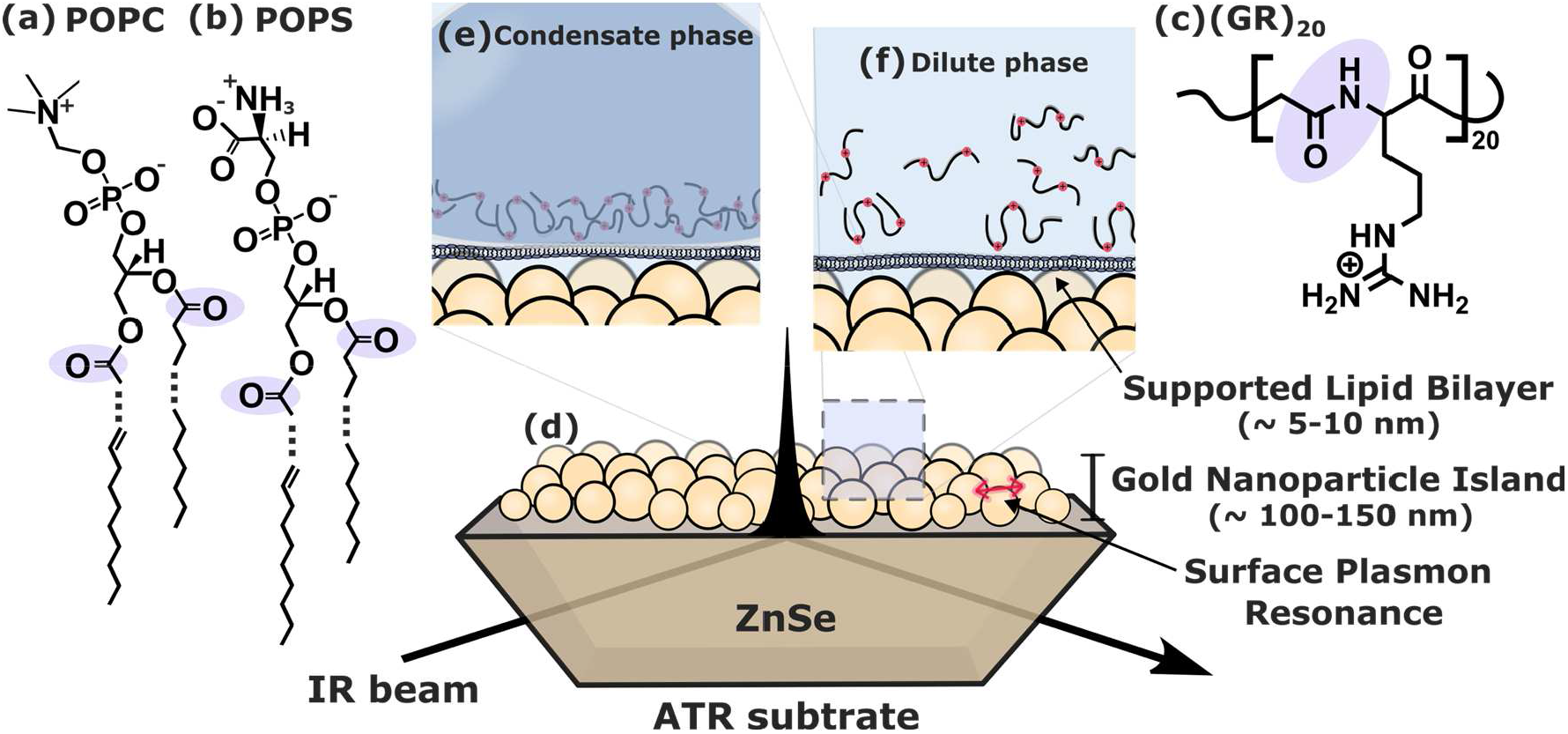
Molecular structure of **(a)** 1-palmitoyl-2-oleoyl-glycero-3-phosphocholine (POPC) **(b)** 1-palmitoyl-2-oleoyl-sn-glycero-3-phospho-L-serine (POPS) **(c)** (GR)_20_ **(d)** Diagram of the surface-enhanced infrared absorption spectroscopy (SEIRAS) setup. An infrared (IR) beam is directed through a high-refractive-index attenuated total reflectance (ATR) substrate; here a ZnSe crystal was used. The resulting evanescent electromagnetic field penetrates the surface and is amplified by the surface plasmon resonance on the gold nanoparticle islands deposited on top of the crystal. The magnified panel illustrates the experimental configuration of the supported lipid bilayer on top of the gold nanoparticles, and (e) condensate droplet interface and (f) dilute phase of (GR)_20_ peptides within the <100 nm probing region. The droplet diameter is ∼3-7 µm, significantly larger than the probed interfacial region.

## Results and Discussion

Phosphatidylserine (PS) constitutes approximately 3-10% of the inner plasma membrane leaflet and is specifically enriched in late secretory and endocytic compartments.^29,30^ Here we explore the effects of 10 mol% POPS addition to in phosphatidylcholine (PC) membranes, where the negatively-charged headgroup induces strong electrostatic interactions with the positively-charged peptides in the condensate. Supported lipid bilayers were formed by vesicle fusion of neutral POPC or POPC/POPS (9:1) mixtures onto gold surfaces, confirmed by carbonyl absorption features near 1730 cm^-1^ (**Figure S1, 2**).^31^ (GR)_20_ condensates, composed of ∼3-7 µm diameter droplets, were prepared in phosphate buffer containing 30% (v/v) PEG to promote phase separation and introduced onto the sample cell prior to measurements. The droplets, initially dispersed in the buffer solution, settled to the surface and reached structural equilibration over a period of ∼200 minutes (*Section S5*). Given the surface-sensitivity of SEIRAS ^24^ the measurements report on the structure of the droplets, in contact with the bilayer within <100 nm of the surface, probing only the structure of peptides at the membrane interface (**Figure 1e**).

Amide I lineshapes directly report on peptide secondary structure. On POPC bilayers, the amide I maximum appeared near 1645 cm^−1^(**Figure 2a**), whereas in POPC/POPS (9:1) bilayers the band is shifted to ∼1660 cm^−1^(**Figure 2b**). Arginine side-chain features at 1585 and 1607 cm^−1^were also observed. The POPC/POPS system also exhibited a negative band near 1730 cm^−1^assigned to the phospholipid ester carbonyl stretches, indicating lipid perturbation through electrostatic interactions. The decreased intensity in the carbonyl region suggests local membrane reorganization or curvature, implying stronger interactions between the condensate and the lipid interface. In addition to SEIRAS, transmission FTIR spectroscopy was used to measure the secondary structure of peptides within the “bulk” condensate droplets (**Figure S7**). For comparison, we also analyzed the dilute peptide phase both in bulk and in the presence of membranes (**Figure 1f, S3**).These reference spectra are in good agreement with recently published spectra of the same systems which examined the changes in secondary structure between the dilute and condensate phase in bulk.^17^ Additional control experiments include SEIRAS spectra of supported lipid bilayers with 30% (v/v) with PEG-only as well as a measurement of (GR)_20_ condensates in the absence of lipid membranes. These control experiments are described in *Section S3*.

**Figure 2.**
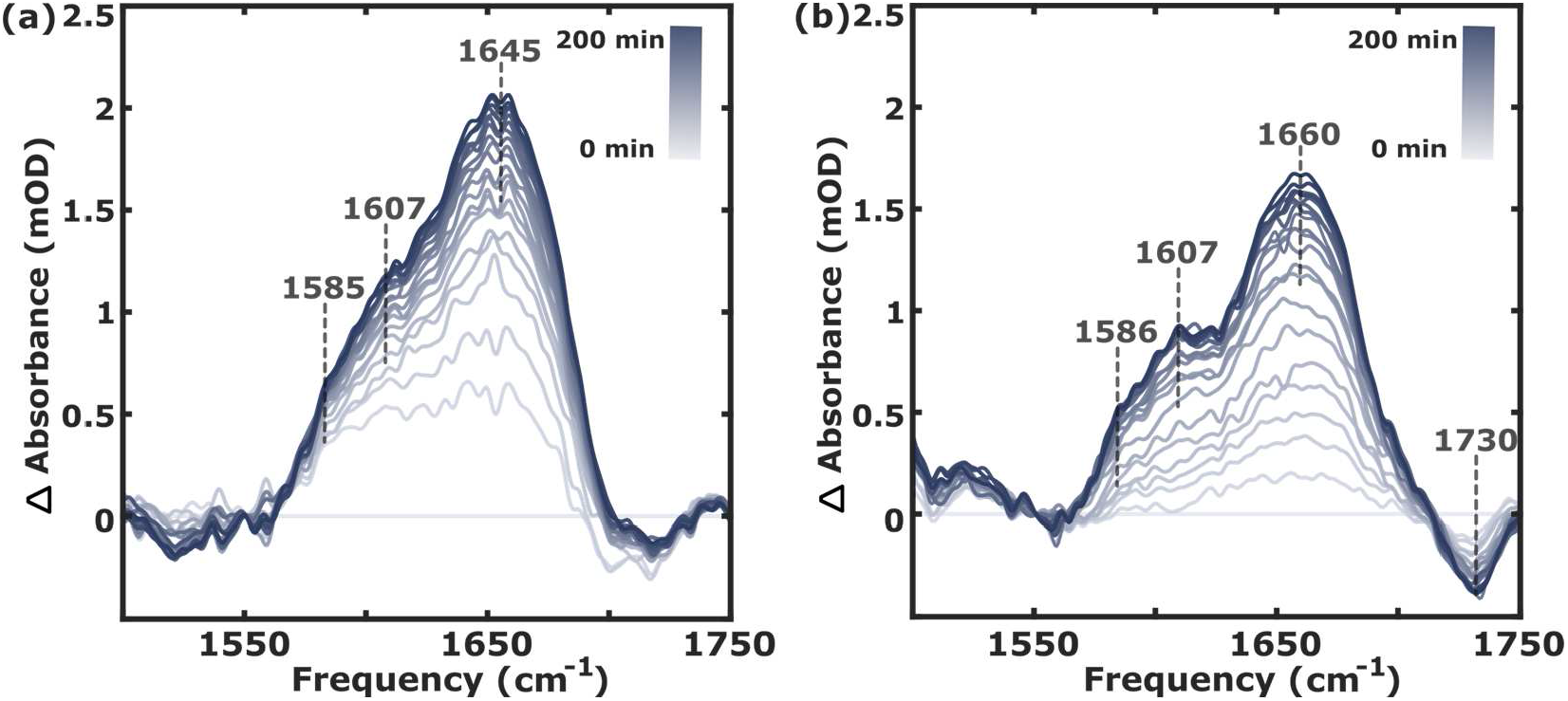
**(a)** Real-time measurements of SEIRA spectra of the arginine side-chain CN stretch (1585-1605 cm^-1^) and amide I region (1550–1750 cm^−1^) of (GR)_20_ condensate phase on the POPC membrane system **(b)** POPC/POPS membrane system, measurement recorded every 2 min over 200 min. The initial (0 minute) spectrum has been subtracted and used as the baseline. The time is indicated in the color bars.

The spectra for both condensate and dilute phase (**Figure S8**) were described by a sum of three Gaussian functions representing β-sheet (1615-1638 cm^-1^), α-helix and disordered (1639-1660 cm^-1^), and β-turn/loop (1653-1691 cm^-1^) structures (**Figure 3a, S12)**.^19^ Based on these assignments, spectra were fitted using a Markov Chain Monte Carlo (MCMC) sampler (*Section S6)*. The α-helix and disordered features, while distinct in larger proteins, are partially overlapped and difficult to distinguish in the present measurements. Considering the peptide’s characteristics as a low-complexity domain (LCD),^32^ and based on previous studies of poly-GR,^33^ this spectral feature is primarily assigned to the disordered conformation. Higher-resolution methods, such as surface-enhanced two-dimensional (2D) infrared spectroscopy with improved signal-to-noise ratios, could further refine the structural assignment in future works.^34^

**Figure 3.**
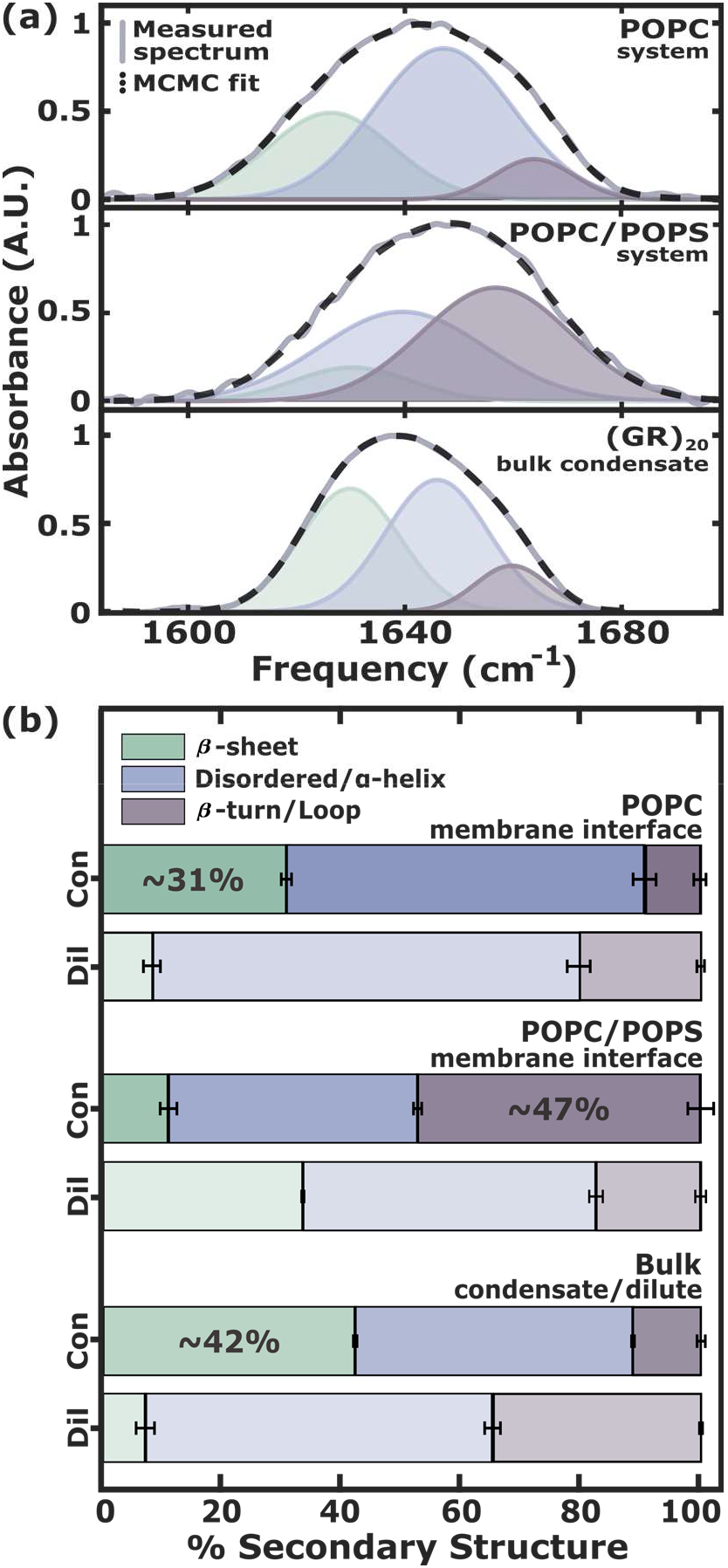
Secondary structure analysis of (GR)_20_ **(a)** Measured spectrum (solid lines) along with Gaussian fits (dashed lines) of the amide I band for the condensate phase. Each Gaussian component is shown underneath the lineshape. **(b)** Relative populations of secondary structures obtained from the spectral fits for each system. Data for the condensate phase are depicted using dark colors, while the corresponding dilute phase results are shown in lighter colors. Error bars were extracted from the standard deviation value from the MCMC sampling distribution.

Transmission FTIR spectra of the bulk condensate droplet, exhibited bands at 1637, 1657, and 1675 cm^−1^, with β-sheets comprising ∼42% of the population (**Figure 3b**). This strong β-sheet signal is in good agreement with previous findings that molecular crowding enhances inter-strand coupling in (GR)_20_, and is further supported by Alphafold3 structural predictions, of the peptide structure.^17^ The disordered band at 1657 cm^−1^was blue-shifted relative to typical values observed in proteins (1639–1654 cm^−1^), likely reflecting partial dehydration under crowded conditions.^35–37^

In contact with the POPC bilayer, Gaussian centers appear around 1632 cm^−1^(β-sheet), 1658 cm^−1^ (α-helix/disordered), and 1679 cm^−1^(β-turn/loop) respectively. The β-sheet fraction decreased to ∼31%, while disordered content increased to ∼60%. The absence of changes in the carbonyl feature around 1724 cm^-1^ indicates minimal bilayer perturbation (*Section S2*), consistent with weak condensate affinity and retention of droplet-like organization. The lineshapes suggest that condensates maintain bulk-like structure at neutral interfaces, with modest redistribution of secondary structure populations. The increased disordered contribution may reflect greater conformational freedom at the boundary, yet the lack of interfacial signatures indicates relatively weak coupling between the condensate and the POPC bilayer.

In the negatively-charged POPC/POPS bilayer system, amide I components appeared at 1639, 1651, and 1672 cm^−1^, accompanied by a further β-sheet reduction and a substantial increase in β-turn/loop (∼47%), indicating significant interfacial structural remodeling induced by the charged lipids. The β-sheet band was blue-shifted relative to POPC, consistent with weaker hydrogen bonding within the condensate and at the protein–solvent interface. These spectral shifts can be attributed to interfacial reorganization under electrostatic interactions similar to conditions to those previously associated with wetting at charged membranes.^6,38^ Indeed, membrane wetting reduces interfacial water reorientation and enhances dehydration,^6,7^ which can induce conformational heterogeneity within the condensate. Molecular dynamics simulations further demonstrate that POPS-containing membranes exhibit enhanced wetting and altered interfacial hydration compared with neutral bilayers, strengthening electrostatic coupling with positively charged peptides.^39^ These effects likely drive the observed redistribution toward β-turn/loop rich structures. Previous work on (GR)_20_ condensates reported loop formation around negatively charged poly-adenosine chains,^17^ suggesting similar loop-like conformations may develop around the phosphate and carboxylate groups of POPS. The concurrent negative band at 1730 cm^−1^ supports membrane reorganization or curvature induced by strong condensate–membrane coupling.

To evaluate whether the observed interfacial reorganization is specific to the condensed state or stems from single peptide-lipid interactions, we measured the dilute phase in the presence of membranes.

Across the bulk solution, neutral POPC, and anionic POPC/POPS conditions, the dilute phase consistently exhibited a high proportion of disordered/α-helix structure. This shared feature reflects the intrinsically disordered character of (GR)_20_ in dilute solution. On the neutral POPC bilayer, the secondary structure profile closely resembled that of the bulk dilute peptide, maintaining the dominant disordered state with a slightly higher proportion of turn structures compared to β-sheets, indicating minimal structural perturbation at the neutral interface. In contrast, the negatively charged POPC/POPS membrane recruited the dilute peptide and induced distinct structural shifts. The relative population ratios of the dilute phase on POPC/POPS were similar to those of the PEG-induced bulk condensate. This similarity indicates that strong electrostatic attraction to the anionic membrane significantly increases the interfacial peptide concentration, suggesting that this 2D interfacial crowding effectively mimics the local peptide density of the droplet. The 1730 cm^-1^ negative band, associated with lipid reorganization, was also present in this charged dilute system (**Figure S3b**), further supporting this strong membrane-peptide interaction and local enrichment at the interface. However, despite these interactions, the recruited dilute peptide did not exhibit the substantial β-turn/loop populations found in the condensate at the POPC/POPS interface. This distinct difference reveals that the unique β-turn-rich architecture is not driven solely by basic electrostatic interactions or local crowding. Rather, the condensate interacts with the anionic membrane as a dense, dynamically cross-linked network driven by multivalent interactions. When this dense condensate droplet binds to the charged lipid bilayer, the spatial constraints restrict extended conformations,^40^ forcing the peptide backbones to adopt the compact chain conformations characteristic of β-turns or loops.

Together, these results show that the interplay between membrane surface charge and the dynamically cross-linked condensate network governs interfacial organization. Neutral POPC bilayers preserve the bulk-like structural profile with minimal perturbation, whereas anionic POPC/POPS membranes induce a pronounced redistribution of secondary structures and local lipid rearrangement. The observed spectral changes reveal that the balance between internal peptide-peptide cohesion and external peptide-lipid adhesion imposes spatial constraints, directly driving the increased β-turn/loop formation at the boundary. These findings connect changes in protein secondary structure to the physical properties of the membrane interface, demonstrating how membrane environments actively direct the molecular architecture of phase-separated condensates.

## Conclusions and Outlook

This study demonstrates that lipid membranes modulate peptide structure within biomolecular condensates. SEIRAS combined with Markov Chain Monte Carlo (MCMC) based analysis shows that variations in membrane composition alter the conformational balance within (GR)_20_ condensates, producing distinct, lipid-dependent secondary structure distributions. Crucially, our comparison with the dilute phase reveals that this interfacial remodeling is uniquely dictated by the cross-linked network of the condensate. The spatial constraints within this condensed network, rather than simple interfacial crowding, drive the distinct structural adaptations observed at the membrane boundary. These results provide molecular level evidence that condensate structure is responsive to the physical and chemical properties of surrounding membranes. By connecting molecular conformation with interfacial organization, this work establishes a framework for understanding how biological interfaces regulate condensate structure, and interactions.

Changes in protein structure may also influence the recruitment of biomolecules to the condensate-membrane interface.^14,41^ While previous studies have shown how condensates reshape membranes, for example by inducing membrane curvature^10^ or forming lipid domains that stabilize the condensate,^9^ our findings demonstrate how membranes, in turn, reshape condensates. The structural organization of condensates is closely tied to their biological functions, emphasizing the interdependence of membrane-condensate coupling. PS lipids act as key regulators of this interaction by tuning interfacial charge and hydration to control protein organization at the surface.^42^ The present results provide a molecular basis for understanding how PS-mediated effects alter condensate structure, with implications for processes such as synaptic assembly and vesicle trafficking.^43,44^ Overall, this work underscores lipid membranes as active participants in condensate biology and opens new avenues for studying the coupling mechanisms of membrane-associated biomolecular condensates.

## Methods

Detailed experimental methods are provided in *Section S1*. Surface-enhanced infrared absorption spectroscopy (SEIRAS) measurements were performed using a Bruker INVENIO spectrometer equipped with a VeeMAX III ATR accessory and an MCT detector. Gold nanoparticle surface for SEIRAS was prepared on ZnSe crystal via electroless deposition. Unilamellar vesicles of POPC and a POPC/POPS (9:1 molar ratio) mixture were prepared in phosphate-buffered saline (PBS, pH 7.4) via freeze-thaw cycling and extrusion (200 nm pore size). (GR)_20_ peptides were purified via acid lyophilization to remove residual trifluoroacetic acid (TFA), and phase separation (500 µM peptide) was induced in 1× PBS using 30% (v/v) PEG 300 as a crowding agent. In addition to these preparative procedures, the Supporting Information contains transmission FTIR data of the (GR)_20_ condensate and dilute phases, the arginine feature subtraction methodology, and a detailed description of the MCMC and Gaussian fitting algorithms. No unexpected or unusually high safety hazards were encountered during the course of this study.

## Acknowledgments

This work was supported by the National Institutes of Health (R35GM133359), and the Welch Foundation (F-1891).

