## Supporting Information for "Lipid Composition Modulates Secondary Structure at the Biocondensate-Membrane Interface"

*Section S1. Sample Preparation*

Infrared Spectroscopy

Surface-enhanced infrared absorption (SEIRA) spectra measurements were carried out using a Bruker INVENIO spectrometer equipped with a Pike Technologies VeeMAX III ATR accessory and a liquid-nitrogen-cooled mercury cadmium telluride (MCT) detector. Spectra were collected at a resolution of 1 $cm^{-1}$, with 128 interferograms averaged for each measurement.

Gold Surface Preparation

Gold surface for SEIRA spectra measurements was prepared on a ZnSe crystal using an electroless gold deposition method as described previously.[1] The ZnSe crystal was pre-heated on a hot plate at 50 °C for 2 min. A 200 μL aliquot of 25 mM $HAuCl_4$ solution in Milli-Q water was applied to the crystal surface for 1 min to initiate gold deposition. Following deposition, the crystals were rinsed thoroughly with Milli-Q water and dried under a gentle stream of nitrogen gas before use.

Lipid Vesicles Preparation

Two lipid vesicle systems were prepared:

1. POPC vesicles: A 25 mg/mL solution of 1-Palmitoyl-2-Oleoyl-sn-Glycero-3-Phosphocholine (POPC) in chloroform (200 μL total volume) was prepared (Avanti Polar Lipids, Alabaster,AL, USA).
2. POPC/POPS (9:1) molar ratio vesicles: A mixed lipid solution with a 9:1 molar ratio of POPC to 1-Palmitoyl-2-Oleoyl-sn-Glycero-3-Phospho-L-Serine (POPS) was prepared by combining 156 μL of 25 mg/mL POPC stock in chloroform with 44 μL of 10 mg/mL POPS stock in chloroform (Avanti Polar Lipids, Alabaster,AL, USA), yielding a total lipid volume of 200 μL at the desired molar ratio.

Lipid solutions were dried under a gentle stream of nitrogen gas for >3 h to form a thin lipid film, ensuring complete solvent removal. A 10× phosphate-buffered saline (PBS) stock solution was prepared by mixing 385 μL of 1 M $KH_2PO_4$ with 615 μL of 1 M $K_2HPO_4$, both dissolved in $D_2O$, following procedures used in previous studies.[2] Given the $D_2O$ environment, the pH meter reading was adjusted to 7.4. The dried films were hydrated with 100 μL of 0.1× PBS buffer and vortexed briefly at a concentration of 50 mg/mL. The suspensions were sonicated for 20 min, followed by 10 freeze–thaw cycles until optically clear, indicating uniform unilamellar vesicle formation. The vesicles were extruded 20 times through a polycarbonate membrane with 200 nm pore size using an extruder to obtain unilamellar vesicles.

$(GR)_{20}$ Condensates

$(GR)_{20}$ peptide (Genscript, USA) was purified by acid lyophilization in $D_2O$ containing 1% DCl to remove residual trifluoroacetic acid (TFA) from synthesis. Condensates were prepared by diluting $(GR)_{20}$ to a final concentration of 500 μM in 1× PBS containing 30% (v/v) polyethylene glycol (PEG) 300 as a macromolecular crowding agent to induce phase separation. Droplet diameter vary from 1~7 μm as reported previously.[2]

*Section S2. Surface Enhanced Infrared Spectroscopy (SEIRAS)-Supported lipid bilayer formation*

SEIRAS selectively enhances vibrational modes with transition dipole moments oriented perpendicular to the metal surface, a property that enables sensitive monitoring of molecular orientation and film formation at interfaces.[3] This feature was utilized to track the formation of supported lipid bilayers. For measurements, vesicles were diluted to one-fourth of their original concentration immediately before being transferred into the ATR chamber, where they fused onto the gold surface to form a supported lipid bilayer. Bilayer formation reached equilibrium after ~2 h and was monitored in real-time by tracking the carbonyl stretching peak (1720-1730 cm$^{-1}$) and the $CH_2$ symmetric and asymmetric stretch region near 2900 cm$^{-1}$ in SEIRA spectra (**Figure S1, S2**).[4] Spectra no longer changed after approximately 120 minutes, as monitored for 3-hour period. The steady signal increase confirmed proper bilayer formation and stability of the supported lipid bilayer. After the bilayer formation, the buffer was replaced with fresh 0.1× PBS buffer, and no change in band intensity or position was observed, ensuring that the bilayer remained intact.

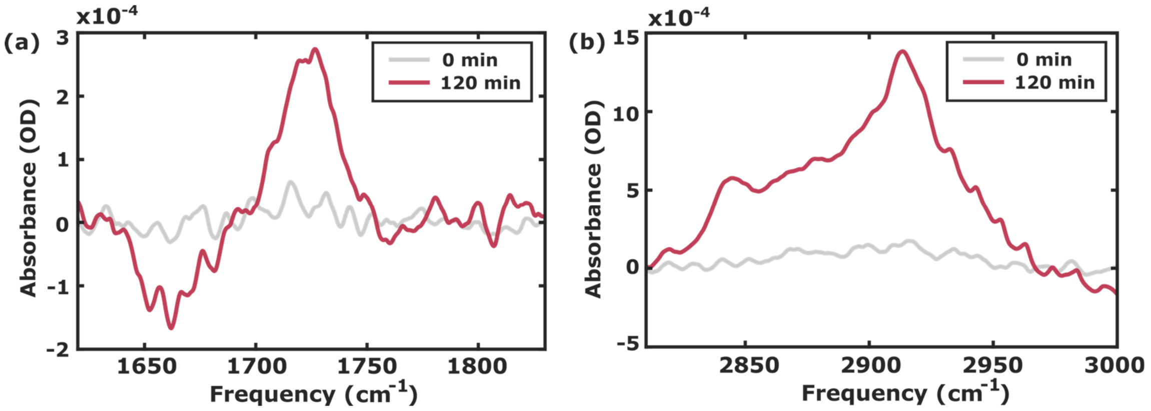


**Figure S1.** SEIRA spectra tracking the lipid bilayer formation of POPC 100% (a). carbonyl (1724 cm$^{-1}$) region of the lipid head group (b) symmetric (2844 cm$^{-1}$) and asymmetric (2913 cm$^{-1}$) stretch mode of $CH_2$ of the acyl chain. The negative feature around 1670 cm$^{-1}$ corresponds to the bending mode of water and represents the displacement of water from the gold layer.

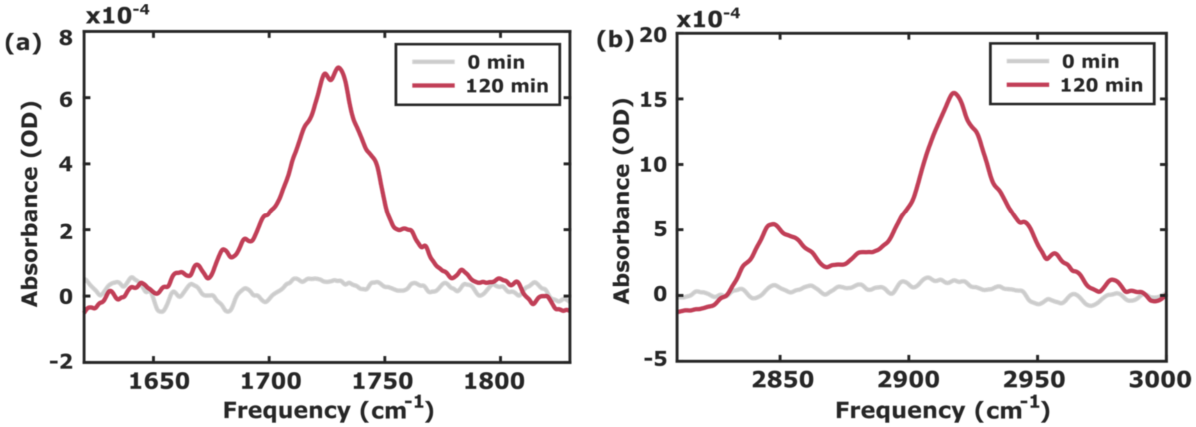


**Figure S2.** SEIRA spectra tracking the lipid bilayer formation of POPC/PCPS (9:1) (a). carbonyl (1730 $cm^{-1}$) region of the lipid head group (b) symmetric (2846 $cm^{-1}$) and asymmetric (2917 $cm^{-1}$) stretch mode of $CH_2$ acyl chain.

*Section S3. Surface Enhanced Infrared Spectroscopy (SEIRAS)-Control experiments*

*Supported lipid bilayer with dilute $(GR)_{20}$ peptide*

After formation and rinsing of the supported lipid bilayer, a 500 μM $(GR)_{20}$ peptide in 1× PBS buffer prepared in $D_2O$ was introduced into the chamber, and SEIRA spectra were collected every 2 minutes. For the 100% POPC membrane (**Figure S3a**), the amide I band gradually increased in intensity over time, reaching ~1.4 mOD with a maximum near 1650 $cm^{-1}$. In the POPC/PCPS (9:1) system (**Figure S3b**), the amide I band appeared around 1658 $cm^{-1}$, and increased to ~0.5 mOD after 200 minutes, accompanied by a smaller decrease (~0.23 mOD) in the negative feature corresponding to the carbonyl region of the lipid headgroup. This gradual rise and distinct peak positions, compared with the condensate phase (**Figure 2**), reflect slower adsorption and a different interfacial configuration of the peptides at the membrane surface. The slower kinetic growth and reduced intensity in the dilute phase, relative to the condensate phase, indicate limited interfacial accumulation of dispersed peptides. This behavior supports that condensates are driven toward the supported bilayer primarily due to their higher density compared to the surrounding buffer.

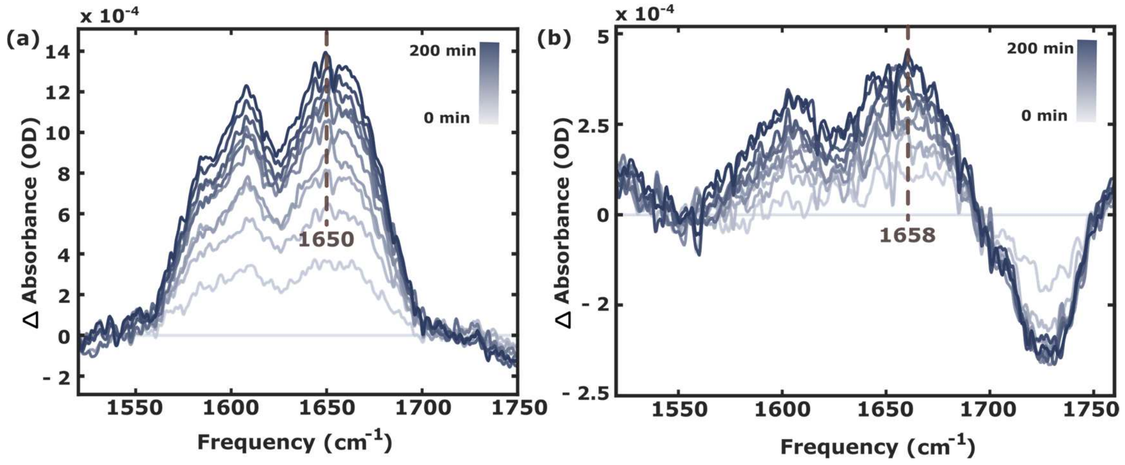


**Figure S3.** Real-time SEIRA spectra of the arginine side-chain feature (1585-1605 $cm^{-1}$) and the amide I region (1550-1700 $cm^{-1}$) of dilute $(GR)_{20}$ peptide with **(a)** POPC 100% membrane and **(b)** POPC/PCPS (9:1) membrane collected over 200 min. The 0 min spectrum was subtracted as a background, and the time is indicated in the color bars.

*Supported lipid bilayer with PEG*

After formation and rinsing of the supported lipid bilayer, 30% (v/v) PEG300 in 1× PBS buffer prepared in $D_2O$ was introduced into the chamber, and SEIRA spectra were collected every 2 min. For the 100% POPC membrane, the C-O stretching band of PEG near 1100 $cm^{-1}$ increased over time, accompanied by a negative feature at ~1725 $cm^{-1}$ corresponding to the C=O stretch of the POPC ester group, with an intensity change of approximately 0.13 mOD (**Figure S4**). The POPC/POPS (9:1) bilayer exhibited a similar trend but with a slightly larger decrease of ~0.24 mOD (**Figure S5**). The difference likely arises from variations in lipid stability on the gold surface.[5] In comparison, the $(GR)_{20}$ condensate system showed a more pronounced negative feature of ~0.5 mOD (**Figure 1b**), indicating that the observed intensity decrease primarily arises from condensate–membrane interactions rather than from PEG alone. These results suggest that while PEG may weakly interact with the membrane, it does not dominate peptide-lipid interactions and instead functions mainly as a macromolecular crowding agent.

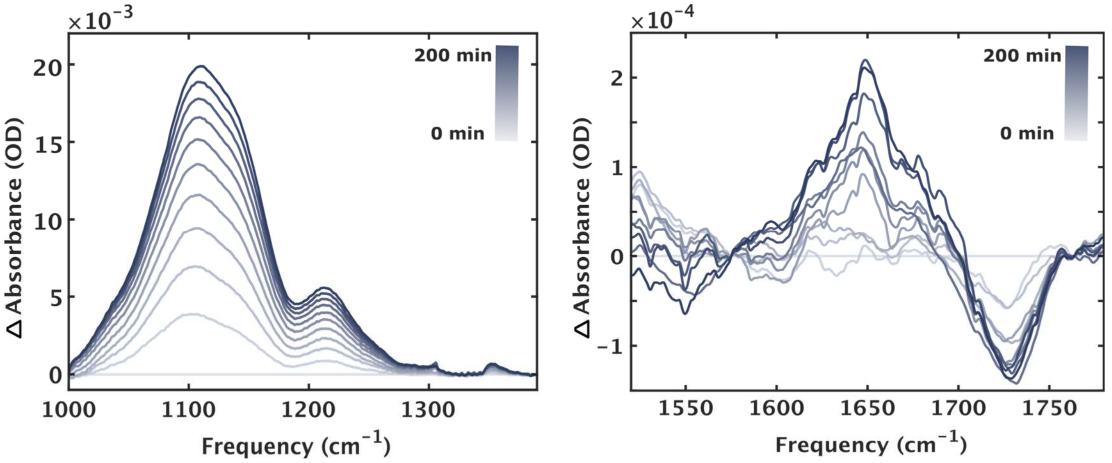


**Figure S4.** Real-time SEIRA spectra of the 30% (v/v) PEG in 1x PBS buffer with POPC 100% membrane collected over 200 min. The 0 min spectrum was subtracted as a background. **(left)** The 1000-1400 $cm^{-1}$ corresponds to C-O stretching vibrations of PEG. **(right)** The 1400-1800 $cm^{-1}$ region includes the carbonyl stretch of the lipid head group intensity decrease around 1725 $cm^{-1}$. The 0 min spectrum was subtracted as a background, and the time is indicated in the color bars.

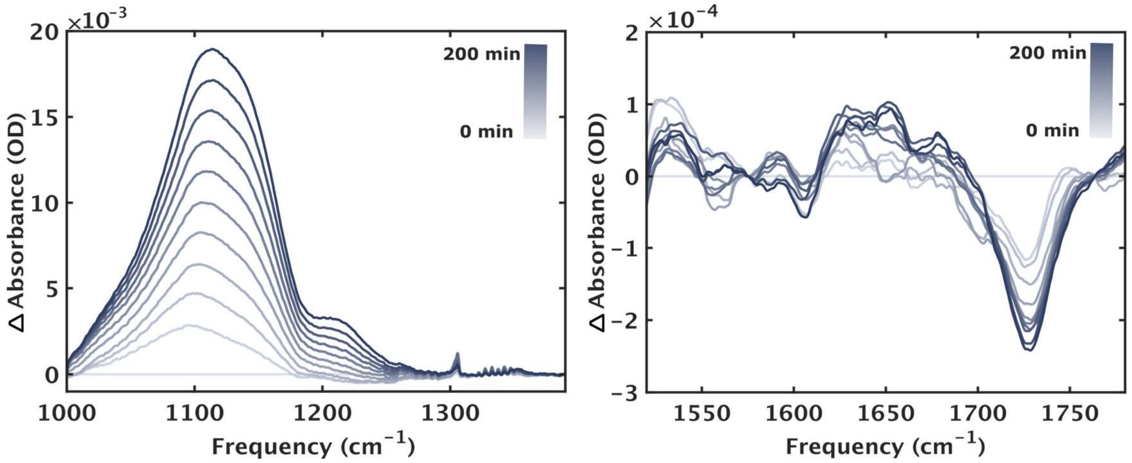


**Figure S5.** Real-time SEIRA spectra of the 30% (v/v) PEG in 1x PBS buffer with POPC/PCPS (9:1) membrane collected over 200 min. The 0 min spectrum was subtracted as a background. **(left)** The 1000-1400 $cm^{-1}$ corresponds to C-O stretching vibrations of PEG. **(right)** The 1400-1800 $cm^{-1}$ region includes the carbonyl stretch of the lipid head group around 1728 $cm^{-1}$. The 0 min spectrum was subtracted as a background, and the time is indicated in the color bars.

*$(GR)_{20}$ condensate without membranes*

Without any lipids, 500 μM $(GR)_{20}$ condensate with 30% (v/v) PEG in 1× PBS buffer prepared in $D_2O$ was introduced into the chamber, and SEIRA spectra were collected every 2 min. The amide I signal populates after 120 min, coinciding with water depletion from the surface, whereas PEG absorption increases continuously from the start. These results indicate that the condensate droplet gradually settle onto the gold surface, but the molecular picture differs markedly from that observed in the presence of a membrane.

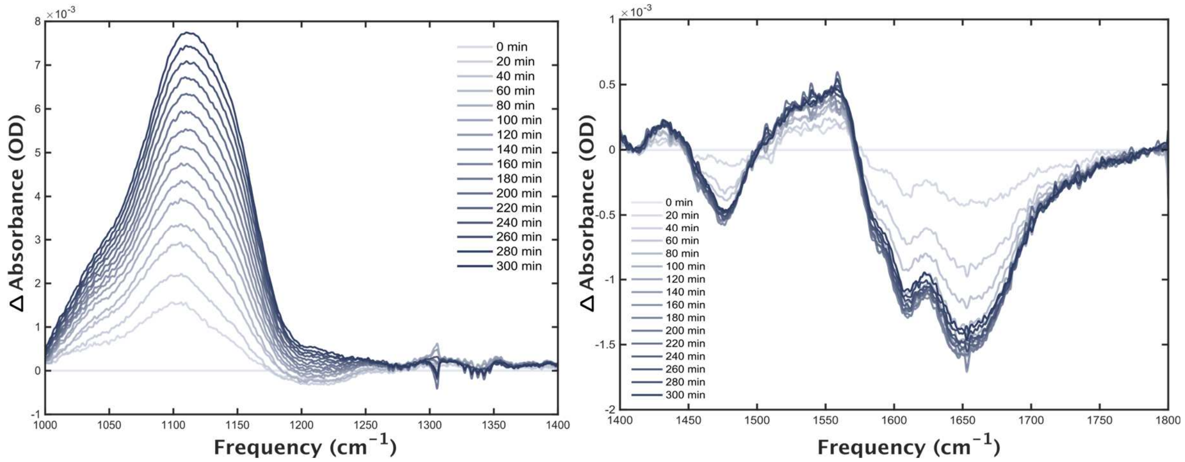


**Figure S6.** Real-time SEIRA spectra of $(GR)_{20}$ condensates without membranes, collected over 300 min. **(left)** The 1000-1400 $cm^{-1}$ corresponds to C-O stretching vibrations of PEG. **(right)** The 1400-1800 $cm^{-1}$ region includes the amide I band of $(GR)_{20}$ condensate. The 0 min spectrum was subtracted as a background.

*Section S4. Gaussian Peak Fitting and Arginine Subtraction*

Gaussian peak fitting was performed by defining an initial model with five peak centers, selected based on reported values for arginine side-chain and amide I vibrations.[6,7] A best-fit model derived from a representative buffer spectrum provided the initial parameter set for subsequent fittings, constrained to variations around these values. The method was applied to all SEIRA spectra as well as transmission FTIR spectra of $(GR)_{20}$ condensates and dilute phase (**Figure S7**). For each spectrum, multiple random initial guesses for amplitudes, widths, and center positions were generated. Amplitudes were initialized between 0 and 1, center frequencies were allowed to fluctuate within ± 2 $cm^{-1}$ of the reference positions (1585, 1605, 1635, 1650, and 1675 $cm^{-1}$), and Gaussian widths varied from 0 to 20 $cm^{-1}$. To minimize sensitivity to starting conditions, each dataset was refitted 50 times with randomized initial parameters drawn from a standard normal distribution within the bounds ("bootstrap fitting"), and the final parameters were calculated as the mean ± standard deviation across all repeats. This procedure ensured reproducibility and stable convergence across all spectra. Gaussian components corresponding to arginine side-chain vibrations were then subtracted from the SEIRA spectra, and the resulting amide I profiles for both condensate and dilute phase are shown in **Figure S8**, with all fitted arginine side-chain parameters summarized in **Table S1** and **S2**.

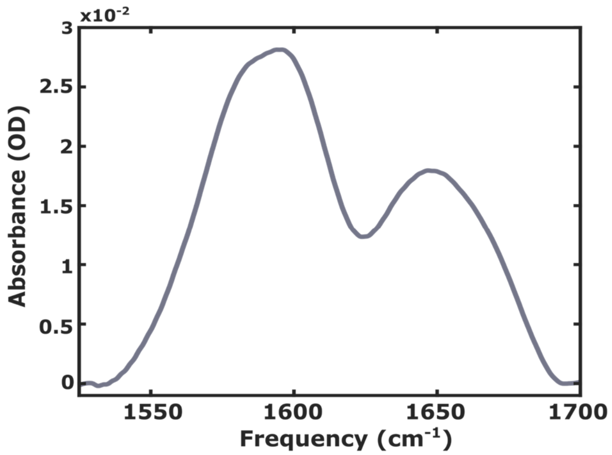


**Figure S7**. Amide I region **(left)** $(GR)_{20}$ condensate **(right)** dilute $(GR)_{20}$ FTIR transmission mode spectrum. The features in the 1550 to 1610 $cm^{-1}$ range corresponds to the side-chain CN vibrations. The band centered around 1650 $cm^{-1}$ corresponds to the amide I region (backbone carbonyl stretches).

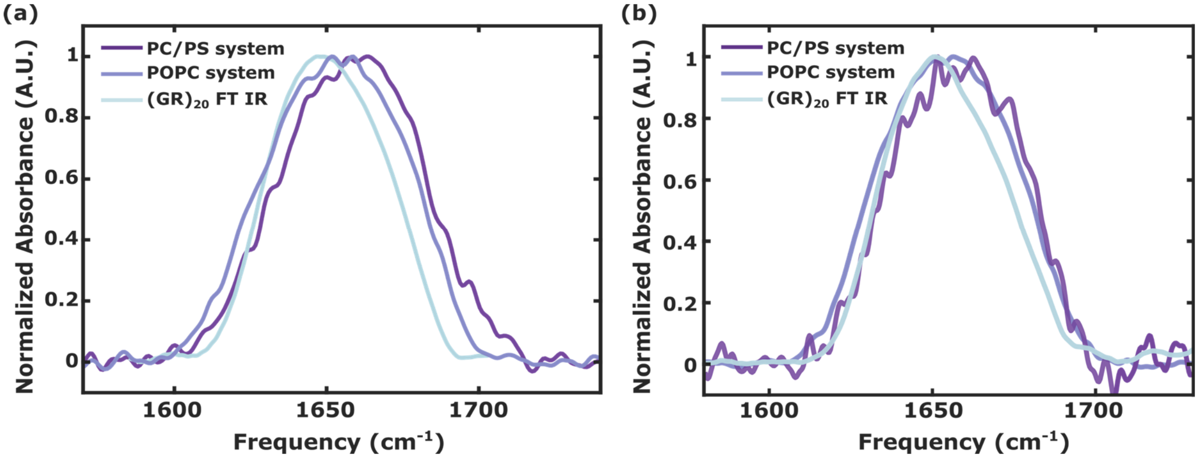


**Figure S8**. Asymmetric and symmetric stretch of arginine sidechain features subtracted SEIRA spectra of $(GR)_{20}$ peptides in **(a)** condensate phase and **(b)** dilute phase.

**Table S1**. Parameters of best fit of bootstrap method (center, amplitude, width) of arginine side-chain. (condensate phase)

| | Amplitude 1 (OD) | Amplitude 2 (OD) | Center freq 1 ($cm^{-1}$) | Center freq 2 ($cm^{-1}$) | Width 1 ($cm^{-1}$) | Width 2 ($cm^{-1}$) |
|---|---|---|---|---|---|---|
| POPC | $0.16 \pm 3.0\times10^{-4}$ | $0.51 \pm 1.0\times10^{-4}$ | $1583 \pm 7.5\times 10^{-3}$ | $1608 \pm 1.5\times 10^{-3}$ | $9.55 \pm 5.0\times10^{-3}$ | $15.04 \pm 0.012$ |
| POPC/POPS | $0.19 \pm 4.4\times10^{-5}$ | $0.48 \pm 3.4\times10^{-4}$ | $1585 \pm 0.38$ | $1608 \pm 0.29$ | $6.15 \pm 0.45$ | $12.02 \pm 0.74$ |
| FTIR | $0.51 \pm 3.6\times10^{-5}$ | $0.71 \pm 7.3\times10^{-4}$ | $1582 \pm 0.05$ | $1605 \pm 0.05$ | $9.53 \pm 0.024$ | $10.1 \pm 0.039$ |

**Table S2**. Parameters of best fit of bootstrap method (center, amplitude, width) of arginine side-chain. (dilute phase)

| | Amplitude 1 (OD) | Amplitude 2 (OD) | Center freq 1 ($cm^{-1}$) | Center freq 2 ($cm^{-1}$) | Width 1 ($cm^{-1}$) | Width 2 ($cm^{-1}$) |
|---|---|---|---|---|---|---|
| POPC | $0.56 \pm 8.3\times10^{-4}$ | $0.78 \pm 2.3\times10^{-4}$ | $1585 \pm 1.70$ | $1608 \pm 0.29$ | $11.22 \pm 1.9\times10^{-3}$ | $9.78 \pm 2.5\times10^{-3}$ |
| POPC/POPS | $0.32 \pm 5.9\times10^{-4}$ | $0.65 \pm 4.6\times10^{-4}$ | $1582 \pm 3.1\times10^{-3}$ | $1607 \pm 0.041$ | $11.64 \pm 0.008$ | $12.16 \pm 0.0398$ |
| FTIR | $0.77 \pm 8.3\times10^{-4}$ | $0.90 \pm 4.2\times10^{-5}$ | $1585 \pm 2.1\times10^{-3}$ | $1608 \pm 5.7\times10^{-4}$ | $8.57 \pm 1.2\times10^{-3}$ | $9.10 \pm 3.4\times10^{-3}$ |

*Section S5. Timescales of peptide-membrane interactions*

Following the spectral decomposition and isolation of the amide I band, the structural dynamics were quantified by tracking the total secondary structure area over time. The total amide band area at each time point was calculated by summing the analytically integrated areas ($A \cdot \omega\sqrt{2\pi}$), where A and ω are the fitted amplitude and width parameters, respectively) of the three core structural components (1635, 1650, and 1675 $cm^{-1}$). Residual bootstrapping was used to robustly determine standard deviations for the integrated areas, accounting for uncertainty in kinetic traces and spectral fitting errors. For each time point, 50 bootstrap iterations were performed by randomly resampling the fit residuals and adding them to the best-fit model prior to refitting, providing highly robust error bars for the extracted areas. Finally, the time evolution of the integrated amide area was modeled using a first-order exponential equation, $y(t) = a(1 - e^{-kt}) + c$ (**Figure S9**). This allowed for the precise extraction of the kinetic rate constant (k) and the characteristic time constant ($\tau = 1/k$) of the structural transition. Based on the extracted characteristic time constants for the two systems ($\tau$ = 64.3 min and 72.7 min), the structural transitions were determined to be approximately 95% and 94% complete, respectively, at 200 minutes ($t \approx 3.1\tau$ and $t \approx 2.7\tau$). Consequently, data collection was maintained up to this point, and the SEIRA spectra recorded at this near-equilibration stage were utilized to interpret the final secondary structure populations.

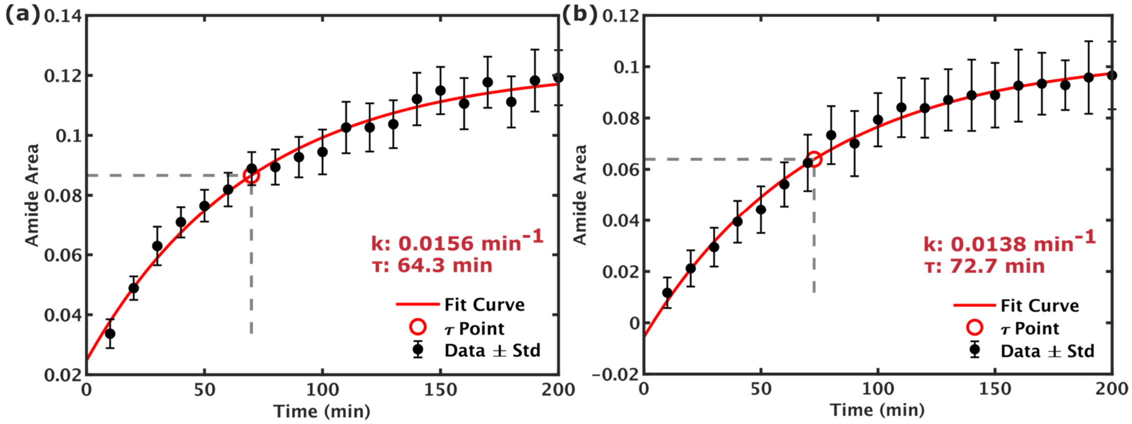


**Figure S9.** The integrated amide I band area (black circles) is plotted as a function of time, with error bars representing the standard deviation from the gaussian fitting bootstrap analysis. A first-order exponential fit (solid red line) to the experimental data yields a structural transition rate constant of τ = 64.3 min and 72.7 min for each $(GR)_{20}$ condensate with **(a)** POPC 100% membrane and **(b)** POPC/PCPS (9:1) membrane. The corresponding τ point (representing 63.2% of the total amplitude change) is highlighted by the red circle and dashed gray lines.

*Section S6. Markov Chain Monte Carlo (MCMC) Gaussian Fitting Method*

Gaussian fitting parameters of the SEIRA spectra were extracted using a Bayesian Markov Chain Monte Carlo (MCMC) approach implemented with the Goodman–Weare affine-invariant ensemble sampler (the "stretch move" algorithm).[8] This sampler evolves an ensemble of walkers in parameter space and is affine invariant, providing improved convergence and mixing for correlated parameters. The spectral model was defined as the sum of three Gaussian components and a noise-scaling term, yielding ten free parameters: three amplitudes ($m_1$–$m_3$), three center frequencies ($m_4$–$m_6$), three widths ($m_7$–$m_9$), and one log-variance parameter ($m_{10}$). The forward model for the absorbance spectrum was expressed as:

$$y_{model}(x;m) = m_1 \exp\left[-\left(\frac{x-m_4}{m_7}\right)^2\right] + m_2 \exp\left[-\left(\frac{x-m_5}{m_8}\right)^2\right] + m_3 \exp\left[-(\frac{x-m_6}{m_9})^2\right]$$

The variance of each data point includes both experimental noise and model-dependent uncertainty:

$$\sigma^2(x;m) = \sigma_{exp}^2 + (y_{model}(x;m)e^{m_{10}})^2 + 10^{-6}$$

Here $\sigma_{exp}$ was estimated from the standard deviation of 20 replicate SEIRA spectra collected near the equilibration time (200 min). In the case of FTIR data, this experimental uncertainty was estimated uniformly across the entire spectrum from the standard deviation at the last 20 data points, representing a peak-free baseline region. The additional noise parameter $m_{10}$ scaled the model uncertainty in log space. The small additive constant ($10^{-6}$) provided as a variance floor to prevent numerical instabilities during likelihood evaluation when $\sigma^2 \rightarrow 0$. Prior bounds were imposed to ensure physically meaningful fits: amplitudes between 0.1-1.0, peak centers between 1600 and 1700 $cm^{-1}$ (with windows applied to each Gaussian to prevent overlap), widths between 9 and 30 $cm^{-1}$, and noise scaling between -10 and 0.1. Walkers were initialized by perturbing parameters obtained from an initial least-squares Gaussian fit. Sampling was performed with 400 walkers for $6\times10^5$ steps, discarding the first 30% as burn-in and thinning by a factor of 30 to reduce autocorrelation. The Goodman–Weare stretch-move acceptance rule was applied to ensure detailed balance. Convergence was assessed by inspecting trace plots and verifying adequate mixing across walkers. Posterior distributions were obtained for all parameters, and the median values and standard deviations were taken as the final Gaussian parameters and their associated uncertainties. Secondary structure populations were computed from the integrated areas of the fitted Gaussian components.

*(GR)$_{20}$ condensate phase*

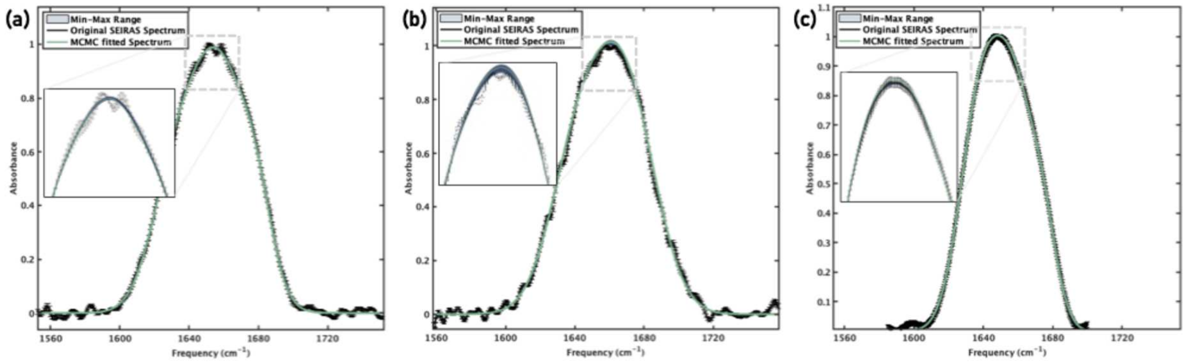


**Figure S10.** The mean posterior spectrum (solid green line) and the 1σ confidence interval (light blue shaded band) illustrate the uncertainty in the fitted spectral profile. The narrow uncertainty region indicates strong convergence and reliable parameter estimation. Each spectra figure shows (GR)$_{20}$ condensate with **(a)** POPC membrane **(b)** POPC/PCPS (9:1) membrane **(c)** without membrane, bulk FTIR data.

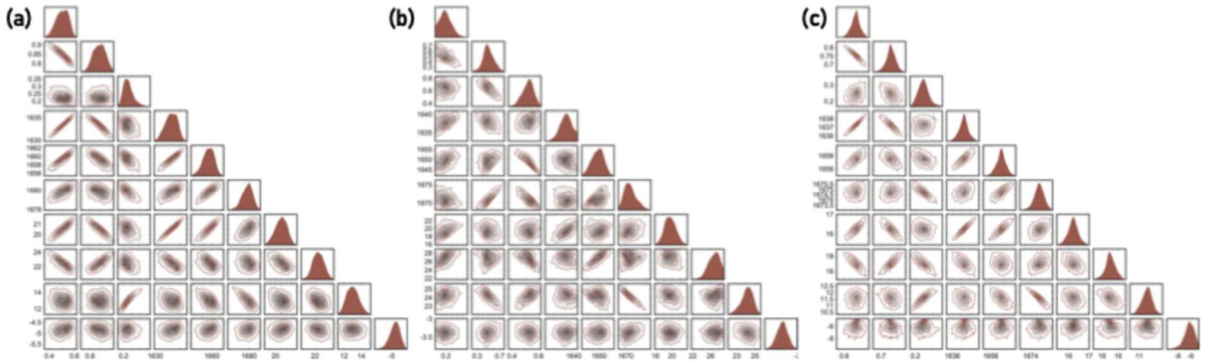


**Figure S11.** Posterior distributions of the fitted parameters were visualized with corner plots showing both marginal distributions (diagonal) and pairwise correlations (off-diagonal). The unimodal and relatively symmetric posterior distributions indicated that the fits were well-constrained and free of strong multimodality. **(a)** POPC membrane **(b)** POPC/PCPS (9:1) membrane **(c)** without membrane, bulk FTIR data.

*(GR)$_{20}$ dilute phase*

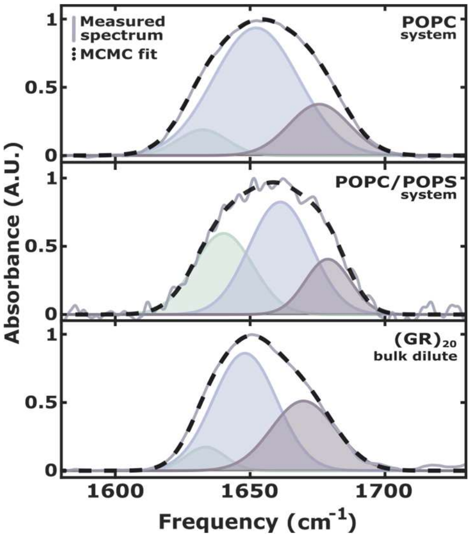


**Figure S12.** Measured spectrum (solid lines) along with Gaussian fits (dashed lines) of the amide I band for the dilute phase for each system. Each Gaussian component corresponds to different peptide secondary structure.

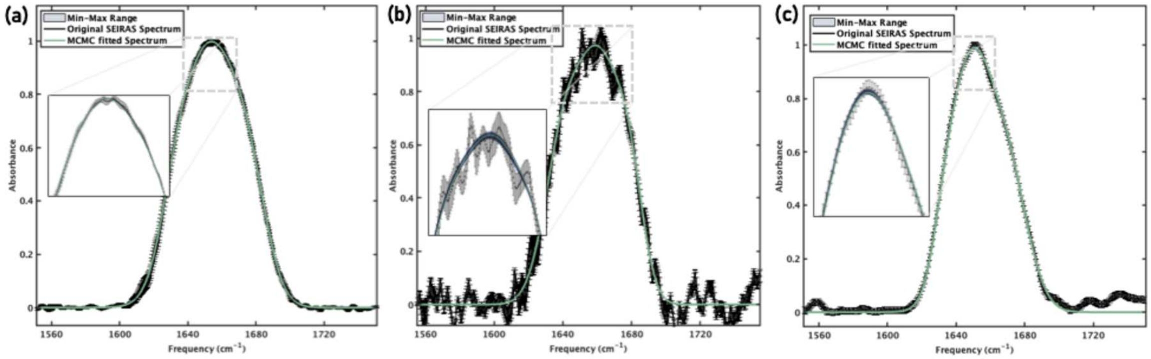


**Figure S13** The mean posterior spectrum (solid green line) and the 1σ confidence interval (light blue shaded band) illustrate the uncertainty in the fitted spectral profile. The narrow uncertainty region indicates strong convergence and reliable parameter estimation. Each spectra figure shows (GR)$_{20}$ condensate with **(a)** POPC membrane **(b)** POPC/PCPS (9:1) membrane **(c)** without membrane, bulk FTIR data.

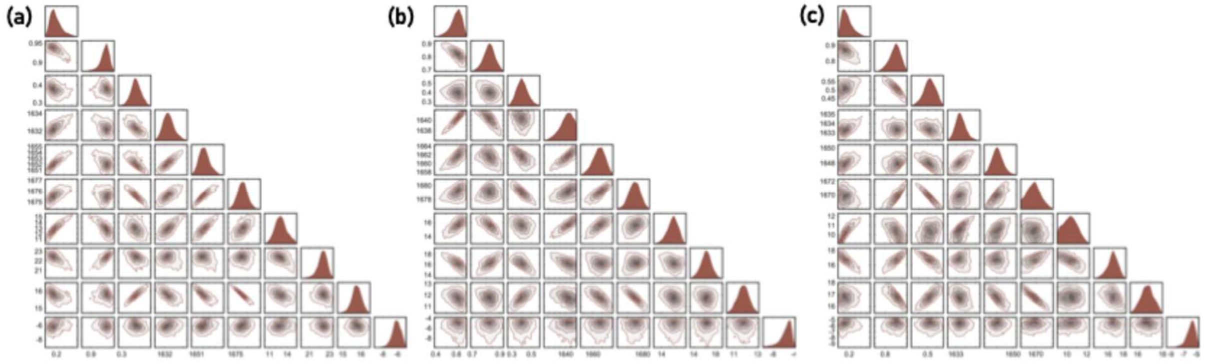


**Figure S14.** Posterior distributions of the fitted parameters were visualized with corner plots showing both marginal distributions (diagonal) and pairwise correlations (off-diagonal). The unimodal and relatively symmetric posterior distributions indicated that the fits were well-constrained and free of strong multimodality. **(a)** POPC membrane **(b)** POPC/PCPS (9:1) membrane **(c)** without membrane, bulk FTIR data.

**Table S3.** MCMC Gaussian fitting result parameters ($m_1$-$m_{10}$) (condensate phase)

| | $m_1$ (A.U.) | $m_2$ (A.U.) | $m_3$ (A.U.) | $m_4$ ($cm^{-1}$) | $m_5$ ($cm^{-1}$) | $m_6$ ($cm^{-1}$) | $m_7$ ($cm^{-1}$) | $m_8$ ($cm^{-1}$) | $m_9$ ($cm^{-1}$) | $m_{10}$ |
|---|---|---|---|---|---|---|---|---|---|---|
| POPC | 0.44 ± 0.056 | 0.88 ± 0.030 | 0.24 ± 0.045 | 1632 ± 1.4 | 1658 ± 1.6 | 1679 ± 0.68 | 19.8 ± 0.61 | 23.1 ± 0.80 | 13.0 ± 0.93 | -4.94 ± 0.24 |
| PC/PS | 0.22 ± 0.069 | 0.52 ± 0.13 | 0.61 ± 0.11 | 1639 ± 1.9 | 1651 ± 2.9 | 1672 ± 1.9 | 20.1 ± 1.5 | 27.4 ± 1.2 | 23.9 ± 0.79 | -3.35 ± 0.13 |
| FTIR | 0.70 ± 0.035 | 0.75 ± 0.027 | 0.26 ± 0.027 | 1637 ± 0.52 | 1657 ± 0.60 | 1675 ± 0.36 | 16.3 ± 0.23 | 16.7 ± 0.62 | 11.7 ± 0.29 | -7.09 ± 0.82 |

**Table S4.** MCMC Gaussian fitting result parameters ($m_1$-$m_{10}$) (dilute phase)

| | $m_1$ (A.U.) | $m_2$ (A.U.) | $m_3$ (A.U.) | $m_4$ ($cm^{-1}$) | $m_5$ ($cm^{-1}$) | $m_6$ ($cm^{-1}$) | $m_7$ ($cm^{-1}$) | $m_8$ ($cm^{-1}$) | $m_9$ ($cm^{-1}$) | $m_{10}$ |
|---|---|---|---|---|---|---|---|---|---|---|
| POPC | 0.18 ± 0.039 | 0.94 ± 0.012 | 0.37 ± 0.029 | 1632 ± 0.68 | 1652 ± 0.94 | 1676 ± 0.45 | 12.5 ± 1.1 | 22.5 ± 0.47 | 15.7 ± 0.27 | -6.66 ± 0.68 |
| PC/PS | 0.61 ± 0.058 | 0.81 ± 0.050 | 0.41± 0.062 | 1640 ± 1.3 | 1661 ± 1.4 | 1679 ± 0.78 | 15.7± 0.89 | 16.1 ± 1.1 | 11.9 ± 0.48 | -6.46± 1.6 |
| FTIR | 0.15 ± 0.067 | 0.86 ± 0.040 | 0.51 ± 0.038 | 1633 ± 0.63 | 1648 ± 0.79 | 1670 ± 0.96 | 10.2 ± 0.84 | 16.9 ± 0.94 | 16.8 ± 0.51 | -6.53 ± 0.78 |
